# Biodegradable Architected Stents for Endoscopic Internal Drainage

**DOI:** 10.64898/2026.05.08.723751

**Authors:** Parima Phowarasoontorn, Yongbin Ko, Zhansaya Makhambetova, Abdel-Hameed Dabbour, Sungyun Sohn, Wegood M. Awad, Oraib Al-Ketan, Mohamed Ali, Juan S. Barajas-Gamboa, Juan P. Pantoja, Ahmed AlZubaidi, Carlos Abril Vega, Panče Naumov, Nader Masmoudi, John Rodriguez, Matthew Kroh, Khalil B. Ramadi

## Abstract

Postoperative gastric leak after bariatric surgery is a serious complication associated with prolonged treatment, repeated interventions, and substantial morbidity. Endoscopic internal drainage using double pigtail stents is widely adopted. However, current stents, originally designed for biliary use and often based on simple cylindrical geometries, are not optimized for post-bariatric gastric leak anatomy, mechanical support, or fluid drainage. Here, we present BRIDGE (Biodegradable aRchitected Internal DrainaGE), a stent concept integrating triply periodic minimal surface (TPMS) architectures to control mechanical compliance, kink resistance, and drainage performance. Using computational modeling, mechanical testing, and benchtop flow studies, we evaluate TPMS designs and identify volume fraction as a key parameter balancing flexibility, structural integrity, and hydraulic performance. TPMS-integrated designs tolerated a 7.1-fold smaller bend radius than a commercial stent without kinking and achieved up to a 2-fold increase in drainage. We also developed a stereolithography-printable biodegradable resin and fabricated a prototype lattice-integrated stent.

**Teaser:** A biodegradable, 3D-printed stent with an architected lattice design improves flexibility, kink resistance, and abscess drainage while eliminating the need for device removal.

## Introduction

Bariatric surgery is an established treatment for morbid obesity and its associated metabolic and cardiovascular diseases. Among currently available procedures, laparoscopic sleeve gastrectomy (LSG) is one of the most performed because of its efficacy and favorable safety profile [1-3]. Despite continued improvements in operative technique and perioperative care, postoperative gastric leak (GL) remains a serious complication that can lead to abscess formation, prolonged hospitalization, repeated interventions, and substantial morbidity. GL occurs in 1-3% of primary cases and up to 10% in revision procedures [4]. Management of gastric leaks has increasingly shifted toward endoscopic approaches, which offer a less invasive alternative to surgical re-intervention (Figure 1A). These techniques include self-expanding metal stents (SEMS), through-the-scope clips (TTSC), over-the-scope clips (OTSC), tissue sealants, suturing systems, and endoscopic internal drainage (EID) utilizing double pigtail stents (DPS) [5-7]. Among these, endoscopic internal drainage (EID) with double pigtail stents (DPS) has gained broad clinical acceptance because it is generally well tolerated and can achieve high rates of leak resolution, up to 89.5% [1-3].

**Figure 1.**
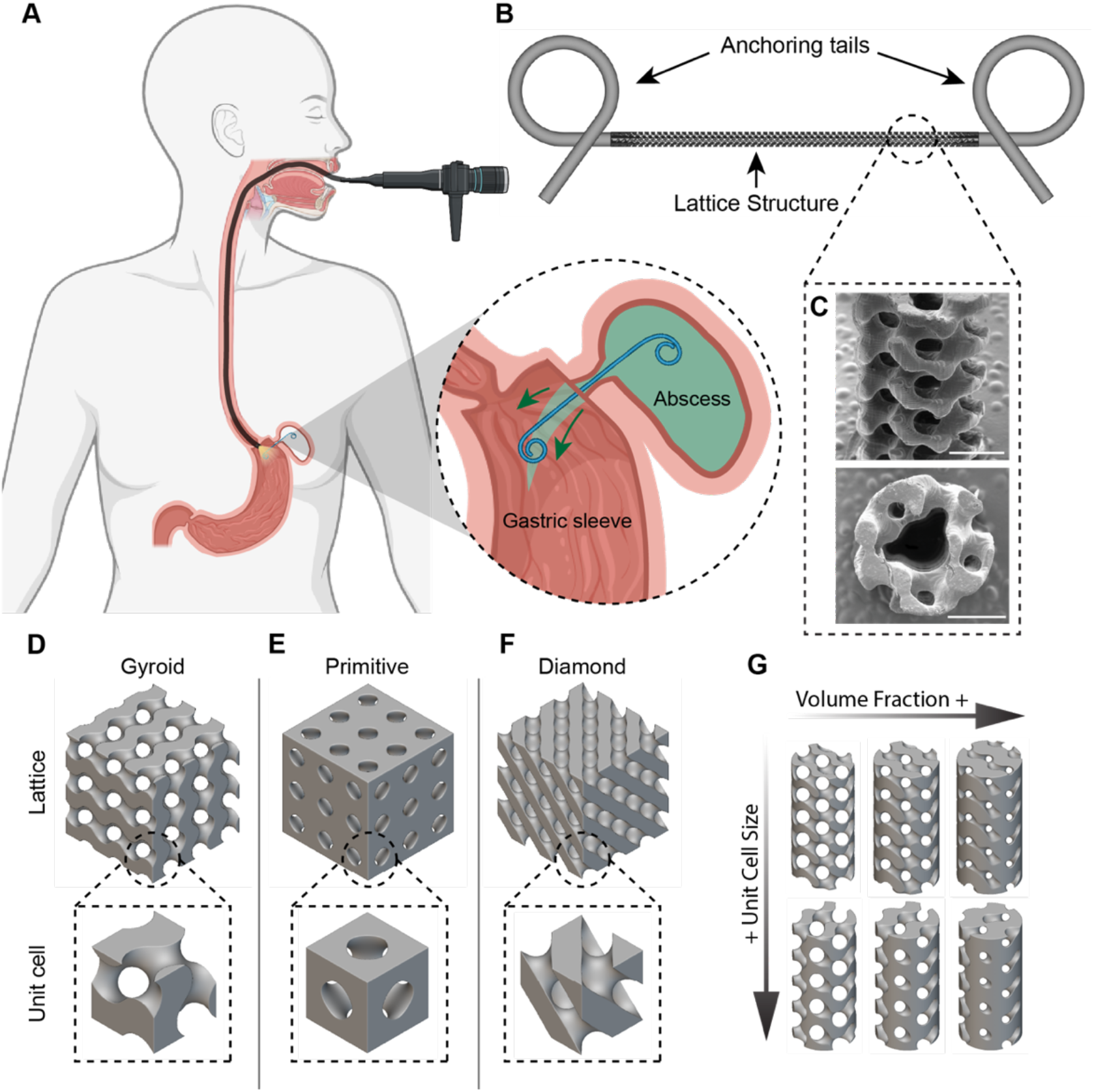
(A) Schematic of the endoscopic procedure used for double-pigtail stent (DPS) deployment. (B) CAD model of BRIDGE. (C) Scanning electron microscopy image of the fabricated 3D-printed prototype (scale bar = 1 mm). (D–F) TPMS topologies evaluated in this work. (G) Representative examples illustrating differences in volume fraction and unit cell size.

However, currently available DPSs were originally developed for biliary applications rather than for the highly variable anatomy of postoperative gastric leaks [8]. In practice, this mismatch creates several important limitations. From an engineering perspective, complex drainage anatomies impose competing requirements on device design. The device must be flexible enough to navigate tortuous or angulated pathways and conform to irregular geometries while remaining sufficiently stable to resist kinking and preserve lumen patency [9, 10]. Second, inappropriate sizing and poor geometric conformity can contribute to stent migration, inadequate anchoring, and insufficient drainage, which may ultimately reduce treatment success and increase the risk of adverse events [9, 10]. Third, because conventional DPSs are typically made from nondegradable polymers, successful treatment still requires a follow-up procedure for stent removal, which contributes to patient burden and procedural cost.

These limitations highlight the need for a gastric leak drainage stent that is not only clinically deployable but also mechanically and hydraulically optimized for this application. Recent reports suggest that, in the management of postoperative gastric leaks, softer and more flexible stent materials may be advantageous for reducing procedure-related tissue and vascular injury compared with standard biliary DPS [2]. Additionally, greater stent compliance could improve navigation and placement within complex postoperative anatomy while providing the anti-kinking behavior required to maintain a patent drainage pathway. Geometric customization enabled by additive manufacturing also offers an opportunity to move beyond conventional extruded circular plastic stents and instead design structures that better balance conformability and flow performance.

In this study, we present BRIDGE (Biodegradable aRchitected Internal DrainaGE) devices, a TPMS (triply periodic minimal surface)-integrated stent concept designed to address key challenges in gastric leak drainage stent through an integrated computational, mechanical, and materials-based approach. We first investigated how TPMS architectures can be used to engineer a stent midsection with tunable flexibility and improved resistance to kink-induced lumen collapse (Figure 1B,C). We next evaluated the effects of TPMS geometry on fluid transport, with the goal of maintaining or improving drainage performance. Finally, we developed an SLA-printable biodegradable resin to enable fabrication of degradable lattice-integrated stents, with the long-term aim of eliminating the need for stent retrieval after abscess resolution. Together, these design strategies establish a proof-of-concept framework for a more compliant, better-draining, and potentially retrieval-free alternative to conventional double pigtail stents for postoperative gastric leak management, with broader applicability to other endoluminal drainage indications.

## Results

### 1. Characterization of TPMS mechanical properties

We first investigated how triply periodic minimal surface (TPMS) lattice topology, unit cell size (UC), and volume fraction (VF) influenced the bending behavior of a tubular lattice structure, which serves as the mid-section of the BRIDGE device (Figure 1D-G). We designed this analysis to identify parameters that could introduce tunable flexibility into the mid-section of a double pigtail stent (DPS) and thereby facilitate deployment in complex anatomy. We distinguished three volume fraction definitions throughout the discussion of the results: designed volume fraction (VF_D_), the target value prescribed during design; nominal volume fraction (VF_N_), the realized TPMS solid fraction measured before central through hole perforation; and effective volume fraction (VF_E_), the realized TPMS solid fraction measured after introducing the central through hole.

We quantified the maximum total displacement produced by a constant bending load across different lattice configurations using parametric finite element analysis (FEA). We fixed both ends of the tube and applied a load of 0.01 N over an area of 3 mm^2^ at the tube midpoint (Figure 2A). For the parametric analysis, we held VF_D_ constant at 0.3 when evaluating the effect of UC and held UC constant at 1 when evaluating the effect of VF. For topology comparison, we evaluated three lattice types (Primitive, Gyroid, Diamond) at VF_D_ of 0.3 and UC of 1. Under these conditions, the Primitive lattice exhibited the lowest maximum displacement (0.87 mm), whereas Gyroid and Diamond lattices showed similar maximum displacements of 2.02 mm and 2.10 mm, respectively (Figure 2B,C). At UC of 1 and VF_D_ of 0.3, the Diamond lattice showed a 33% smaller average feature size than the Gyroid lattice (0.1005 ± 0.0545 mm vs 0.1500 ± 0.0530 mm). We selected the Gyroid architecture for further study because it combined high bending compliance with a larger average feature size than the Diamond lattice at matched design parameters, making it a more practical candidate for fabrication.

**Figure 2.**
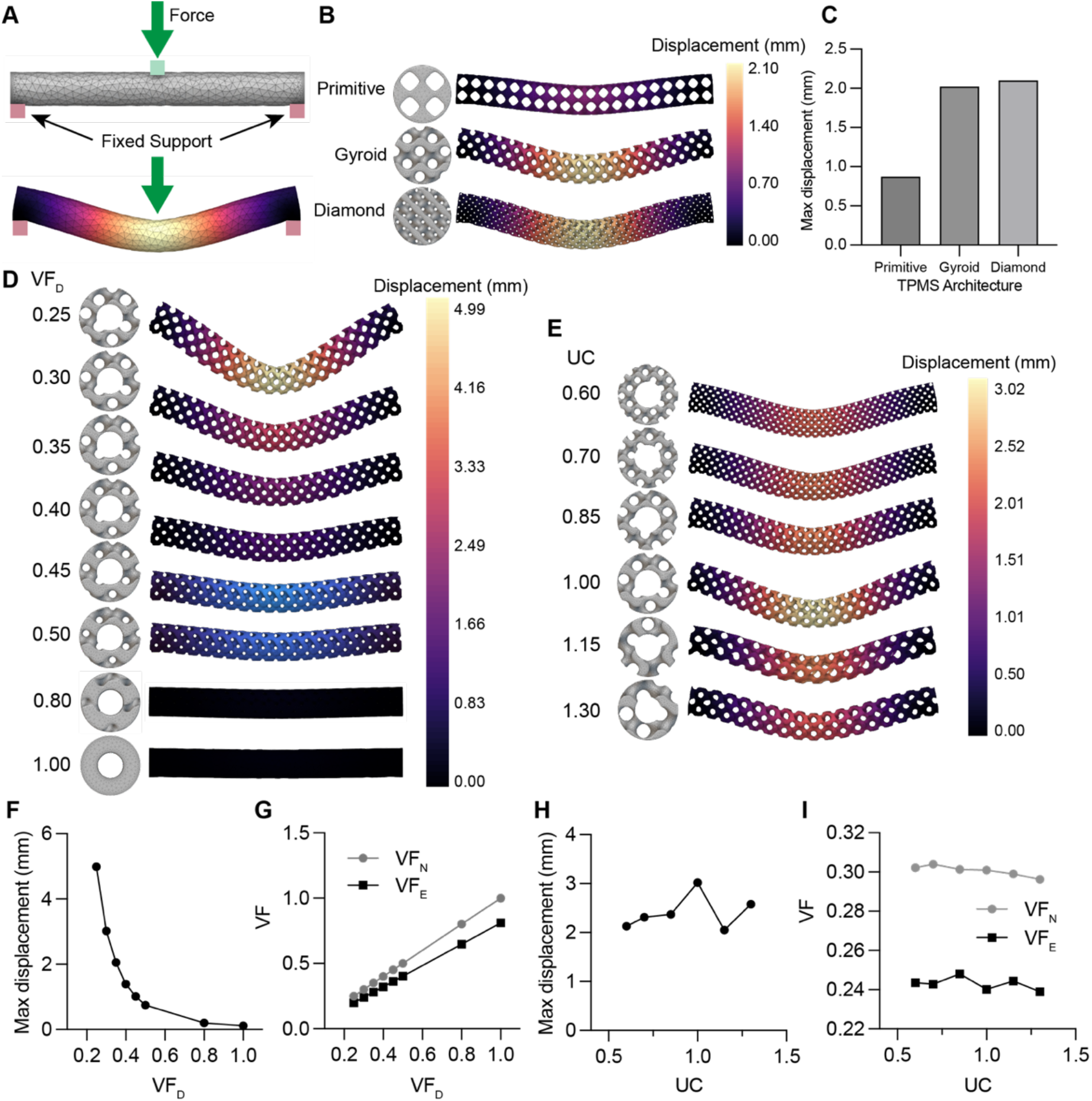
Finite element analysis of TPMS structures under quasi-static bending. (A) Schematic of the applied boundary conditions. (B) Representative deformation results for full TPMS cylinders with Gyroid, Primitive, and Diamond topologies under bending load. (C) Corresponding maximum displacement values for each topology. (D,E) Bending simulation results for Gyroid tubes with a 1 mm inner diameter at varying volume fraction and unit cell sizes. (F) Maximum displacement of Gyroid tubes with fixed UC = 1 and varying VF. (G) Volume fraction before through-hole perforation (VF_N_) and after perforation (VF_E_) for Gyroid UC = 1 with varying VF. (H) Maximum displacement of Gyroid tubes with fixed VF_D_ = 0.3 and varying UC. (I) Volume fraction before through-hole perforation (VF_N_) ad after perforation (VF_E_) for Gyroid VF_D_ = 0.3 with varying UC.

We next extended this analysis to a geometry that more closely resembles the intended DPS design by introducing a 1 mm central through-hole to accommodate standard endoscopic guidewire delivery (Figure 2D,E). Here, we considered a 0.035 inch (0.889 mm) guidewire commonly used for placement of 7 Fr DPS [11]. Using the Gyroid lattice as a representative topology, we first varied VF_D_ from 0.25 to 1 (solid tube) while maintaining UC of 1 and found that increasing VF produced a monotonic decrease in maximum displacement from 4.990 mm to 0.112 mm (97.7% decrease), consistent with increasing bending stiffness (Figure 2F). This trend was consistent with the VF_E_ measured in the design software before and after introducing the through hole (VF_E_ = 0.199 to 0.811, linear fit R^2^ = 1). Although the through hole removed a greater absolute volume at higher VF_D_ (0.051 mm^3^ at lowest VF_D_ vs 0.819 mm^3^ at highest VF_D_), the relationship remained approximately linear over the tested range, consistent with the monotonic reduction in displacement (Figure 2G). In contrast, varying UC from 0.6 to 1.3 at a prescribed VF_D_ of 0.3 produced a non-monotonic displacement response, with the largest maximum displacement observed at UC = 1 (3.02 mm) (Figure 2H). Because VF_D_ was prescribed before introducing the through hole, the perforation altered the VF_E_ and circumferential material distribution differently across UC values depending on the alignment between the TPMS topology and the hole position. This effect was reflected in the post-cut effective VF, which exhibited a local minimum at UC = 1 (VF_E_ = 0.240) and likely contributed to the corresponding peak in displacement (Figure 2I). Together, these results show that both TPMS topology and geometric design parameters can be used to tune tubular bending compliance, while also highlighting that introducing a guidewire lumen can create secondary geometric effects that influence the apparent bending response.

We experimentally evaluated the tensile behavior of perforated 3D-printed TPMS Gyroid tubes across VF and UC design spaces (Figure 3A). All specimens were fabricated from Flexible 80A resin using a Formlabs Form 3+ printer. At a fixed UC = 1, increasing VF_D_ from 0.2 to 0.5 produced an approximately linear increase in ultimate load, from 3.35 ± 0.39 N to 7.34 ± 0.70 N (Figure 3B). Based on the gross tube cross-sectional area of 1.94 mm^2^, we calculated an apparent structural modulus and found that it also increased with VF_D_, from 1.23 ± 0.20 MPa to 2.56 ± 0.36 MPa (Figure 3C). By contrast, varying UC from 0.85 to 2 at fixed VF = 0.3 produced a non-monotonic tensile response, with the highest ultimate load observed at UC = 1 (4.78 ± 0.46 N) and lowest ultimate load of 2.35 ± 0.18 N at UC = 2 (Figure 3D). We observed the local minima in apparent structural modulus was observed at UC = 0.85 and 1.5 (0.64 ± 0.11 MPa and 0.63 ± 0.04 MPa, respectively) (Figure 3E). These data identify VF as a more predictable determinant of tensile strength, whereas UC exerts a nonlinear effect on axial stiffness.

**Figure 3.**
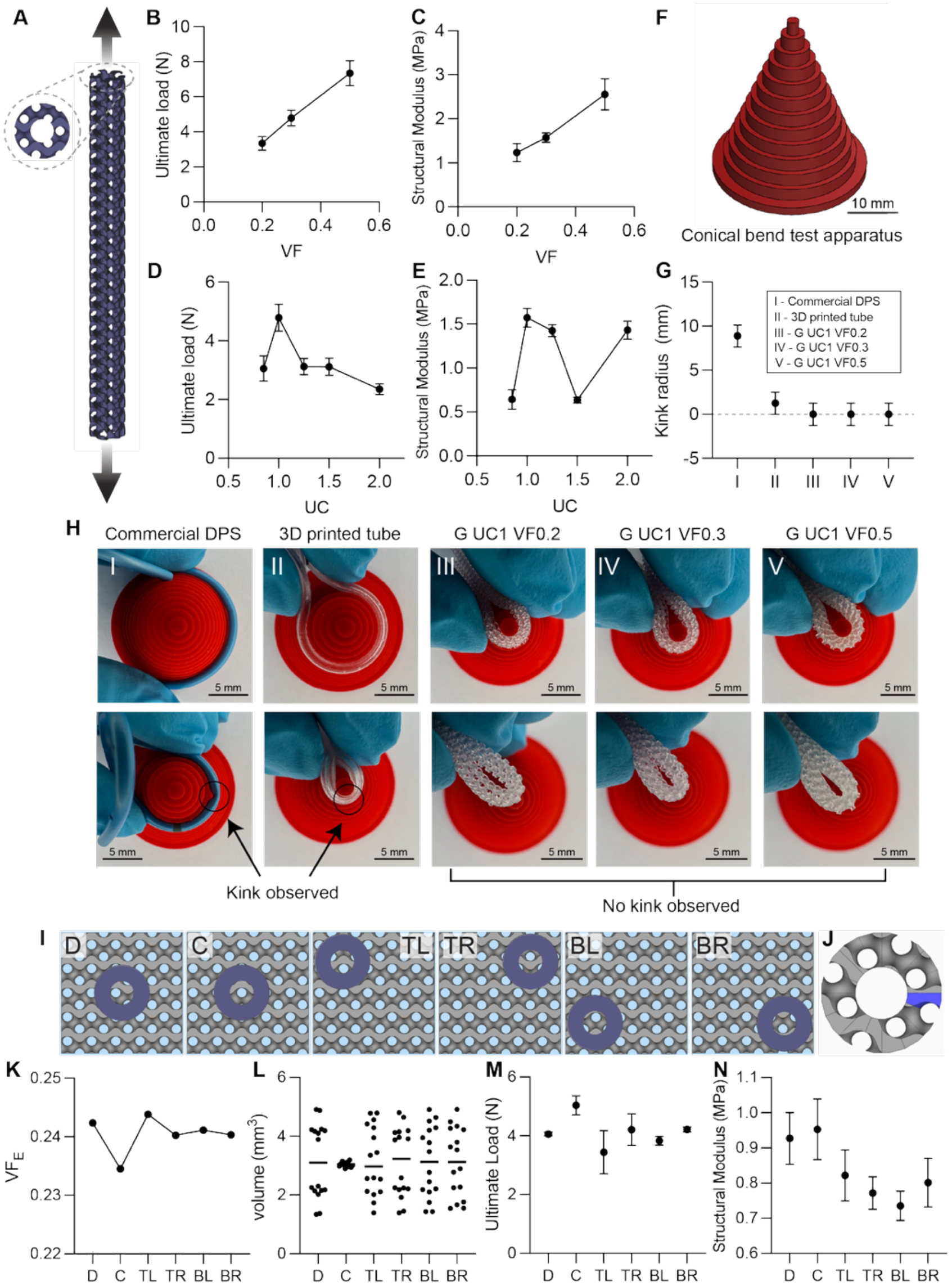
Mechanical characterization of 3D-printed TPMS Gyroid tubes. (A) Schematic of the tensile testing setup. (B,C) Ultimate load and corresponding apparent structural modulus of Gyroid tubes with fixed unit cell size (UC = 1) and varying volume fraction. (D,E) Ultimate load and corresponding apparent structural modulus of Gyroid tubes with fixed volume fraction (VF = 0.3) and varying unit cell size. (F) CAD rendering of the conical bending test apparatus, with diameters decreasing from 30 mm to 1.25 mm in 1.25 mm increments. (G) Maximum kink radius of the commercial polyethylene (PE) stent, 3D-printed Flexible 80A tube, and TPMS Gyroid tubes with UC = 1 and varying volume fraction. (H) Representative images of each specimen before kinking (top) and at the onset of kink formation (bottom), showing minimal lumen collapse in the Gyroid tubes even when folded onto themselves. (I) Six tube positions evaluated for the Gyroid design (UC = 1, VF = 0.3): default (D), center-aligned (C), top left (TL), top right (TR), bottom left (BL), and bottom right (BR). (J) TPMS tube segmented into 16 equal angular sectors for sectional volume analysis. (K) Effective volume fraction (VFE) of the different alignments after through-hole perforation. (L) Sectional volume distribution for each alignment. (M, N) Ultimate load and apparent structural modulus of 3D-printed samples fabricated from Flexible 80A resin.

We then compared the kinking radius of the 3D-printed Gyroid tube with that of a commercial polyethylene 7 Fr Advanix™ biliary DPS (Boston Scientific) used as a clinically relevant benchmark (Figure 3F). Using the ISO 25539-2 definition of kink radius, we determined the minimum curvature at which lumen loss exceeded 50% [12]. The commercial DPS exhibited a kink radius of 8.875 mm, whereas the 3D-printed tube maintained lumen patency to a substantially smaller radius of 1.25 mm (Figure 3G). At a curvature radius of 1.25 mm, the Gyroid tubes showed no measurable lumen loss and retained >90% of their initial lumen diameter even under complete folding; this represents a 7.1-fold improvement in kink resistance over a commercial biliary DPS (Figure 3H). The Gyroid architecture provided markedly improved resistance to kink-induced lumen collapse compared with a clinically used polyethylene DPS.

Finally, we investigated how guidewire-lumen alignment influenced the mechanical performance of the Gyroid architecture at constant UC = 1 and VF_D_ = 0.3. Six tube positions were evaluated: default (D), center-aligned (C), top-left (TL), top-right (TR), bottom-left (BL), and bottom-right (BR) (Figure 3I). To assess whether lumen position also altered material uniformity, each perforated tube was segmented into 16 equal angular sectors and the material volume in each sector was quantified (Figure 3J). Geometric analysis showed that the total VF_E_ after perforation was lowest for the center-aligned configuration (0.235) and highest for the top-left configuration (0.244), indicating that the center-aligned lumen removed the greatest amount of material overall (Figure 3K). The center-aligned configuration exhibited the lowest coefficient of variation in sector-wise volume (2.623%), with a maximum difference of 37.7% relative to the other alignments, indicating the most uniform circumferential material distribution around the lumen (Figure 3L). Despite having the lowest total VF_E_, the center-aligned configuration achieved the highest ultimate load (5.04 ± 0.32 N) and apparent structural modulus (0.953 ± 0.086 MPa) in uniaxial tensile testing (Figure 3M,N). Together, these results indicate that tensile performance depended not only on the total amount of material remaining after perforation but also on how uniformly that material was distributed around the load-bearing cross-section, providing an additional design parameter for optimization of BRIDGE devices.

### 2. Fluid dynamic characterization of TPMS tubes

We performed computational fluid dynamics (CFD) simulations to determine how TPMS topology, VF, and UC influence the fluid transport performance of a BRIDGE device. We modeled a representative TPMS lattice segment of 2 mm in length and 2.3 mm in diameter within a concentric cylindrical domain of 6 mm in length and 3 mm in diameter and prescribed a constant inlet pressure of 668 Pa, consistent with standard adult intra-abdominal pressure, with atmospheric pressure imposed at the outlet (Figure 4A) [13]. We assumed steady-state, laminar flow of a Newtonian fluid under no-slip wall conditions. Prior to CFD, we conservatively calculated Reynolds numbers to be 4.40 to 8.32, validating our laminar flow assumption. Volumetric flow rate was obtained by surface integration of the outlet velocity at the prescribed analysis plane. Representative streamline distributions and cross-sectional velocity maps are shown in Figure 4B and 4C. Under matched conditions, the Gyroid topology achieved a flow rate of 0.151 mL/s, which was 16.2% and 9.4% higher than Diamond (0.130 mL/s) and Primitive (0.138 mL/s), respectively (Figure 4D). At fixed VF_D_ = 0.3, increasing UC from 0.85 to 1.50 increased flow rate from 0.139 to 0.227 mL/s (63.3% increase) (Figure 4E). In contrast, at fixed UC = 1, increasing VF_D_ from 0.2 to 0.5 decreased the flow rate from 0.207 to 0.0985 mL/s (52.4% decrease) (Figure 4F).

**Figure 4.**
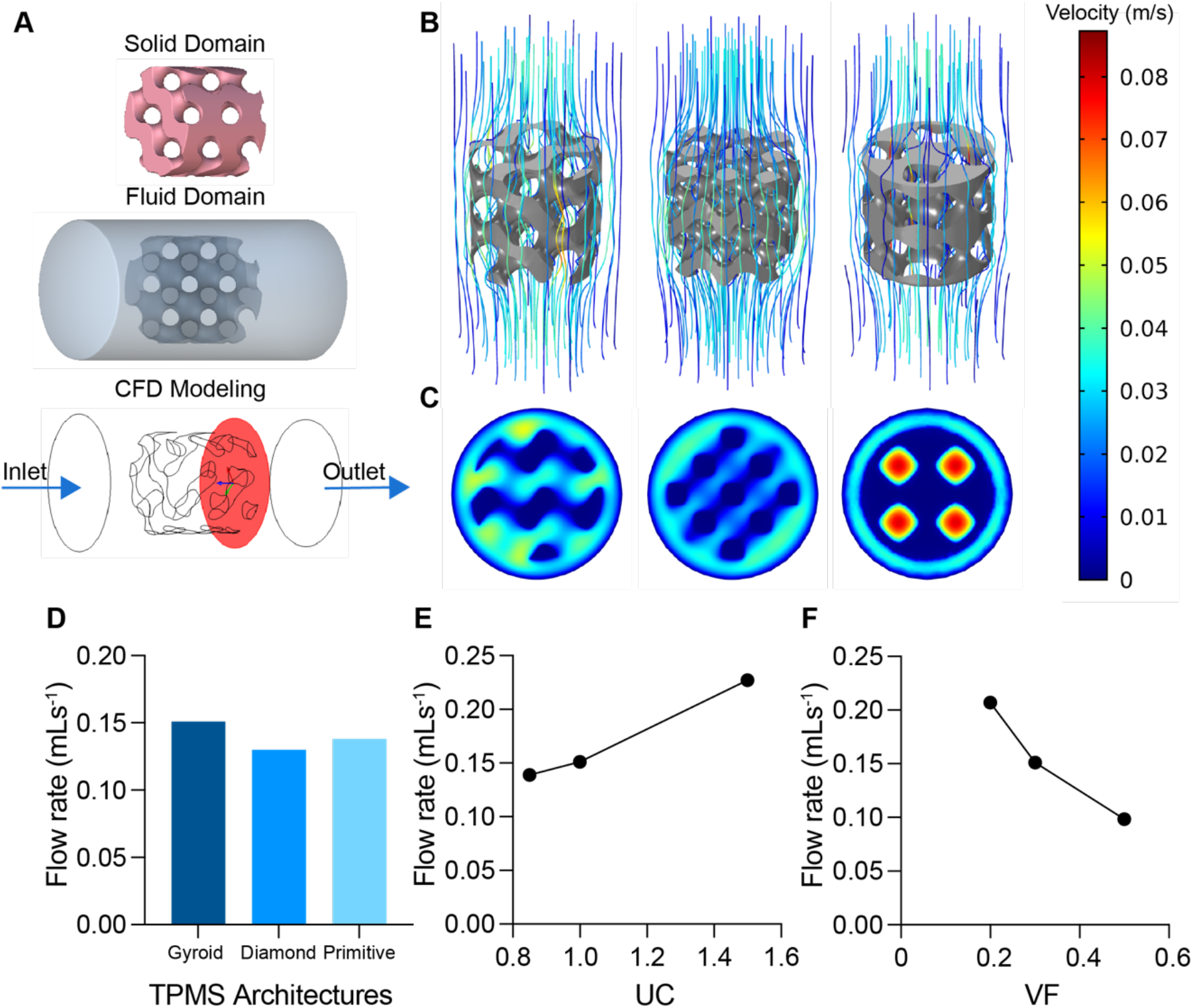
(A) Schematic of the CFD model geometry and boundary conditions. (B,C) Representative flow simulation results for Gyroid, Diamond, and Primitive TPMS topologies. (D) Flow rates of TPMS structures with matched unit cell size (UC = 1) and volume fraction (VF = 0.3) across different topologies. (E) Flow rates of Gyroid structures with fixed volume fraction (VF = 0.3) and varying unit cell size. (F) Flow rates of Gyroid structures with fixed unit cell size (UC = 1) and varying volume fraction.

We then evaluated selected 3D-printed stents in a gastric leak benchtop model by measuring the cumulative mass of drained abscess fluid phantom over time (Figure 5A, B). The UC1VF0.2 and UC1VF0.3 designs achieved flow rates of 41.7 ± 0.4 µL/s and 41.0 ± 0.6 µL/s, respectively, both exceeding that of the commercial control (20.8 ± 0.1 µL/s), whereas the UC1VF0.5 design showed a lower flow rate of 19.2 ± 0.4 µL/s (Figure 5C). These findings confirm that the CFD-predicted trend of decreasing flow rate with increasing VF was consistent with experimental benchtop results, supporting the validity of the simulation as a design-screening tool. Our results also indicate that, at appropriately selected lattice parameters, the TPMS-integrated mid-section can enhance drainage performance relative to a commercial stent while retaining the structural flexibility needed for deployment.

**Figure 5.**
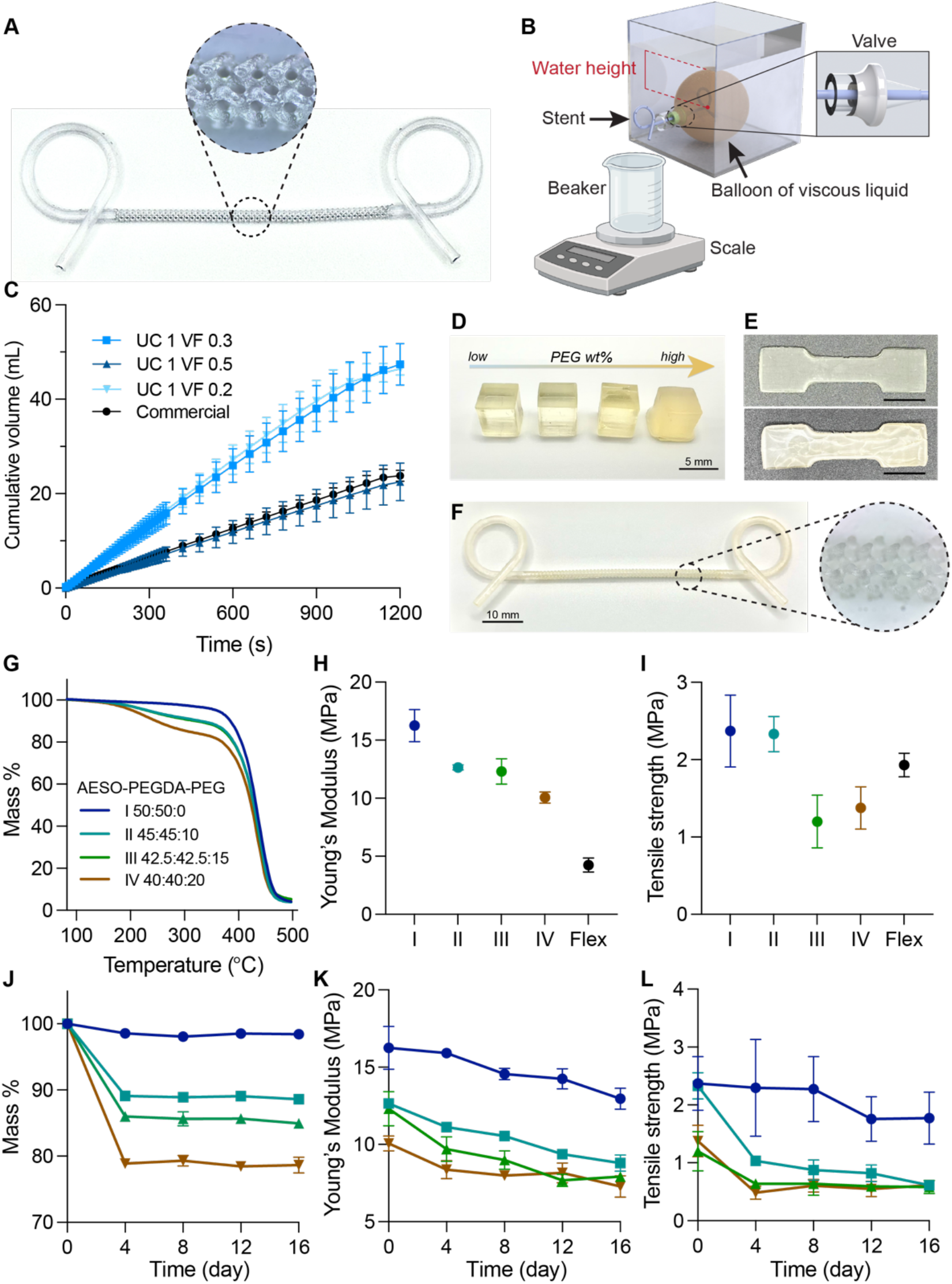
(A) Image of the 3D-printed BRIDGE fabricated from Flexible 80A resin. (B) Experimental fluid dynamics testing platform. (C) Time-dependent cumulative drained volume comparing the commercial DPS and 3D-printed lattice stents (n=3). (D) SLA-printed 50 mm cubes of AESO:PEGDA:PEG resin formulations with increasing PEG content. (E) Representative dog-bone specimens of Type IV resin at day 0 (top) and day 16 (bottom) with a scale bar of 1 cm. (F) SLA-printed BRIDGE fabricated from Type III resin. (G) Thermogravimetric analysis of AESO:PEGDA:PEG resin formulations with varying compositions. (H,I) Young’s modulus and tensile strength of Type I–IV resins compared with Flexible 80A resin prior to degradation testing. (J–L) Percentage mass change, Young’s modulus, and tensile strength of Type I–IV resins following accelerated degradation at 55 °C.

### 3. Biodegradable resin formulation compatible with SLA printing

To develop a biodegradable resin compatible with stereolithography (SLA) printing for BRIGDE fabrication, we evaluated four formulations of acrylated epoxidized soybean oil (AESO) polyethylene glycol diacrylate (PEGDA, M_n_=700), and polyethylene glycol (PEG, M_n_=200) mixtures, with AESO:PEGDA:PEG weight ratios of I (50:50:0), II (45:45:10), III (42.5:42.5:15), and IV (40:40:20) (Figure 5D). Dog-bone specimens were printed for mechanical testing during the degradation study (Figure 5E). Using formulation III as a representative composition for prototype fabrication, we optimized the printing parameters to successfully manufacture a G_UC1_VF0.3 BRIDGE (Figure 5F).

Thermogravimetric analysis (TGA) showed a composition-dependent effect on thermal stability (Figure 5G). All formulations exhibited negligible mass loss up to approximately 150 °C. The onset decomposition temperature (*T*_*onset*_) was approximately 390 °C for all groups. The principal degradation stage occurred between approximately 360 and 480 °C, corresponding to decomposition of the crosslinked polymer network. The temperature at 10% mass loss (T_10%_) was approximately 382 °C, 310 °C, 300 °C, and 235 °C for samples I—IV, respectively, while the temperature of maximum degradation rate (*T*_*max*_) occurred at approximately 440 °C for all samples. Formulation I showed the highest thermal stability, whereas formulation IV showed the earliest degradation onset. All formulations had residual masses of approximately 4% at 500 °C.

We next evaluated the mechanical properties of the printed samples through tensile testing. All specimens were fabricated using identical printing parameters on a customized Form 3+ system, and sample thicknesses were measured at each time point (Table 1). Young’s modulus decreased progressively with increasing PEG content. Compared with Flexible 80A resin, which had an elastic modulus of 1.93 ± 0.15 MPa, the AESO-based formulations exhibited moduli ranging from 1.20 ± 0.34 MPa to 2.37 ± 0.47 MPa (Figure 5H). Tensile strength showed a non-monotonic dependence on PEG composition (Figure 5I).

**Table 1.**
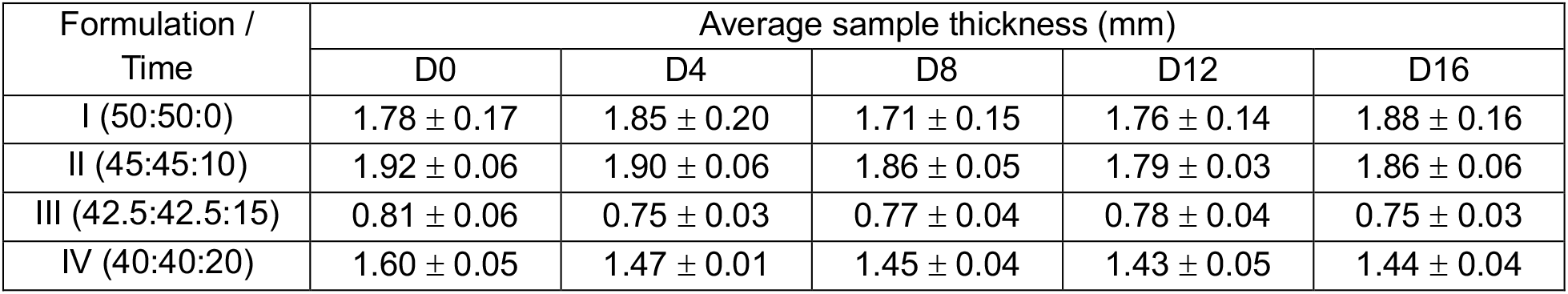
Average 3D-printed dog-bone sample thicknesses.

Accelerated acidic degradation revealed a composition-dependent decline in mass retention and mechanical performance (Figure 5J–L). All formulations showed an initial decrease in mass followed by slower changes at later time points. Formulation I retained the highest mass throughout the study, with final mass retention of 98.4 ± 0.2%, whereas formulations II, III, and IV retained 88.6 ± 0.1%, 84.9 ± 0.7%, and 78.6 ± 1.2%, respectively (Figure 5J). Young’s modulus decreased from 16.3 ± 1.4 to 13.0 ± 0.7 MPa for formulation I (20.2% decrease), 12.7 ± 0.2 to 8.8 ± 0.5 MPa for formulation II (30.7% decrease), 12.3 ± 1.1 to 7.9 ± 0.2 MPa for formulation III (35.8% decrease), and 10.1 ± 0.5 to 7.3 ± 0.7 MPa for formulation IV (27.7% decrease) over the degradation period (Figure 5K). Tensile strength likewise declined from 2.4 ± 0.5 to 1.8 ± 0.5 MPa (25.0% decrease), 2.3 ± 0.2 to 0.6 ± 0.1 MPa (73.9% decrease), 1.2 ± 0.3 to 0.6 ± 0.1 MPa (50.0% decrease), and 1.4 ± 0.3 to 0.6 ± 0.1 MPa (57.1% decrease) for formulations I–IV, respectively (Figure 5L). Overall, higher PEG content was associated with lower mass retention and greater losses of stiffness and strength during acidic degradation.

## Discussion

No commercially available device has been specifically developed for endoscopic internal drainage of gastric leaks, despite the widespread off-label use of double pigtail stents (DPS) for this purpose. Our work demonstrates that integrating a triply periodic minimal surface (TPMS) lattice into the mid-section of a DPS provides an approach to simultaneously improve bending compliance and drainage performance. Current DPSs used for endoscopic internal drainage are largely adapted from biliary applications and are typically manufactured as extruded polymer tubes with limited geometric tunability. Unlike biliary strictures or ducts, post-bariatric leaks often involve irregular extraluminal cavities and tortuous drainage paths, which place different demands on device conformability and patency. Our approach uses mathematically driven lattice architectures as a design variable while preserving the familiar overall DPS form factor, ensuring compatibility with standard endoscopic delivery workflows. This is especially relevant for post-bariatric gastric leaks, where irregular leak tracts, abscess cavities, associated stenosis, and complex postoperative anatomy can make device placement and effective drainage challenging. Our results show that TPMS architecture can be used not only to tailor the mechanical response of the stent mid-section, but also to preserve or enhance flow performance relative to a commercial DPS.

Among the three TPMS topologies evaluated, Gyroid provided the most favorable overall combination of bending compliance, flow performance, and fabrication feasibility. Finite element analysis showed that Gyroid and Diamond exhibited similarly compliant bending responses, both of which were substantially more deformable than Primitive topologies, while Gyroid also maintained a larger average local feature thickness. This larger feature thickness may reduce susceptibility to print defects or local fracture during handling and deployment. This mechanical trend is consistent with prior comparisons of the same TPMS families, in which the Primitive architecture exhibited a relative structural modulus more than twice that of Gyroid and Diamond and deformed through a combination of strut stretching and buckling, whereas Gyroid and Diamond showed more bending-dominated deformation [14, 15]. In this context, the greater compliance of Gyroid and Diamond observed here is likely attributable to both their lower structural stiffness and their greater ability to accommodate the applied load through bending rather than through stretch-dominated deformation. Notably, this interpretation is informed by prior compression-based mechanical studies rather than direct bending experiments [14]. In addition, our FEA treated the printed elastomer as an isotropic, linear elastic material under quasi-static loading, and incorporation of nonlinear, time-dependent, and cyclic loading conditions would yield a more physiologically relevant representation of in vivo deformation.

Gyroid produced the highest flow rate among the three architectures at matched unit-cell size and volume fraction, in CFD simulation. This trend is consistent with prior pore-scale TPMS studies showing that, among comparable porous configurations, Gyroid can exhibit the highest permeability and the lowest pressure drop among topologies, indicating lower overall hydraulic resistance under the same driving conditions [16]. Together, these findings indicate that topology selection should consider mechanical compliance, drainage efficiency, and manufacturability in parallel rather than in isolation.

We used CFD primarily as a comparative design-screening tool, rather than for computation of absolute values. The trend of total flow rate observed in CFD and in the benchtop GL model across varying VF was consistent, with lower VF achieving higher flow rate. The representative-volume CFD model captured the relative influence of topology, VF, and UC on hydraulic performance, but it was simplified to reduce computational demands and therefore involved several assumptions and adjustments. The simulation focused on a section of the connective path between the abscess and the stomach with a full TPMS structure and no guidewire lumen. The experimental system included the full printed device with a through-hole perforation, a one-way 3D-printed valve, and a more realistic drainage path, all of which could contribute to deviations from simulated absolute values. The CFD model also assumed steady, single-phase, incompressible, laminar Newtonian flow and did not capture the full pigtail geometry, cavity heterogeneity, or time-varying pressure conditions during drainage. The present model is a useful screening framework, and could be further developed into a tool for personalized patient-specific simulations of hydraulic behavior by incorporating anatomically realistic leak geometries derived from patient imaging data.

Architected geometries such as TPMS are challenging to optimize even for specific applications, given the broad parameter and design space. We found that volume fraction was the most predictable geometric tuning parameter across both the mechanical and fluid analyses. Increasing VF monotonically decreased simulated maximum displacement under bending load, increased ultimate tensile load, increased apparent structural modulus, and decreased simulated and experimental flow rate. These monotonic trends are advantageous because a predictable design parameter is simpler to optimize. In contrast, the effect of unit cell size was more complex and non-monotonic in TPMS tubes with the introduced clinically necessary 1 mm guidewire lumen. The structural responses appear to depend on the lumen positioning relative to the local TPMS geometry, thereby altering both the effective volume fraction and the radial material distribution. Angular segmentation analysis showed that center-aligned through-hole design had a lower coefficient of variation in sectional volume, indicating a more uniform circumferential material distribution around the lumen. This suggests that the remaining lattice preserved more balanced axial load paths, which may help explain why these specimens achieved higher ultimate load despite having lower effective volume fraction. Tensile strength appeared to depend not only on total remaining material, but also on how evenly that material was distributed after perforation. The tensile modulus reported here is an apparent structural modulus based on the gross tube cross-sectional area rather than the true solid surface area, so these values are best interpreted as comparative design metrics. Future work pairing geometric descriptors with local stress mapping or failure localization would more directly resolve how perforation geometry redistributes load within the lattice. UC is a meaningful but secondary design variable whose effect must be interpreted in the context of the perforated geometry. Additionally, this observation suggests that lattice-lumen alignment and size should be considered design features that require optimization to avoid unintended losses in the structural integrity of BRIDGEs.

Improved stent flexibility is only clinically relevant if the drainage lumen remains patent under deformation. ASTM F3505 defines kink resistance in terms of maintaining lumen patency during bending and recognizes loss of lumen under curvature as the relevant failure mode rather than flexibility alone. Current anti-kinking designs in medical catheters and tubular implants are dominated by reinforcement-based tubular designs, in which kink resistance is achieved by combining flexible polymers with braided metallic layers or coil reinforcement [17-19]. A recent vascular-graft study introduced a coil-reinforced multilayer design specifically to improve kink resistance while acknowledging the inverse tradeoff with flexibility [17]. In contrast, our BRIDGE device embeds anti-kinking ability directly into the drainage-bearing mid-section through a tunable TPMS architecture, rather than adding a separate reinforcing layer, thereby simplifying fabrication while also providing tunable mechanical compliance. This unique characteristic may be especially beneficial during endoscopic deployment and positioning through tortuous or irregular postoperative anatomy, which requires the stent to bend readily without sacrificing drainage function. In addition, recent ureteral stent literature suggests that softer or more flexible stents may improve patient tolerance, supporting the broader rationale for compliant stent designs, provided that lumen patency is maintained under bending [20]. Because kink behavior was assessed under controlled benchtop loading, fatigue testing under repeated physiologically relevant bending cycles is an important next step to determine how well this anti-kinking performance is maintained over time. Ex vivo or in vivo deployment studies would further clarify how the compliant lattice behaves during navigation, positioning, and tissue-constrained bending. Taken together, these observations suggest that a geometry-driven anti-kinking strategy may offer a useful alternative to conventional braid or coil-reinforced designs when both conformability and drainage performance are critical.

Beyond the mechanical and hydraulic advantages demonstrated, the biodegradable material platform offers the possibility of temporary internal drainage without a secondary retrieval procedure. This is clinically-relevant because current protocols often require repeated interventions over several weeks, with each patient undergoing multiple endoscopic sessions. Recent clinical reports indicate that management of post-sleeve gastrectomy leaks commonly requires repeat endoscopic reassessment at 4-6 weeks and often involves approximately three endoscopic procedures over an average treatment course of 57.5 days (8 weeks) [21, 22]. The need to limit prolonged indwelling time is also supported by broader stent literature showing that long-dwell stents are associated with complications such as occlusion, migration, infection, and epithelial hyperplasia that can complicate removal [21]. Hence, a degradable stent with a functional lifetime of 6-8 weeks could be especially attractive, as it coincides with the relevant clinical window for leak resolution and serial endoscopic management.

Our results suggest that our AESO:PEGDA:PEG system provides a promising starting point and could be optimized for gastric drainage applications. We identified SLA printing as a suitable fabrication method because it provides the smooth surface finish and printing resolution needed to manufacture BRIDGEs. However, SLA-printable materials that degrade within a clinically relevant timeframe are not currently available off the shelf. We therefore designed this resin system to balance printability, mechanical integrity, and degradation behavior under acidic conditions relevant to the stomach. AESO primarily contributed to degradability, PEGDA acted as a reactive diluent that reduced viscosity and supported photocrosslinking, and PEG functioned as a non-reactive plasticizing component that increased flexibility while maintaining biocompatibility. This strategy was informed by prior AESO- and AESO:PEGDA-based systems, which have shown favorable printability, cytocompatibility, and degradation behavior in vitro and in vivo, including in bone tissue engineering applications [23-25]. PEG further provided a practical route to tune hydrophilicity and network flexibility, consistent with its broad use in biomedical devices and pharmaceutical formulations [26]. Together, these design choices support our goal of creating a biocompatible, 3D-printable, mechanically flexible material from commercially accessible components without requiring complex synthesis.

Our data show that PEG content strongly influenced the thermal, mechanical, and degradation behavior of the resin network. The TGA results indicated that increasing PEG content reduced thermal stability, while the additional early-stage dip observed at higher PEG content suggests the presence of thermally labile segments or volatile fractions associated with PEG incorporation before breakdown of the primary crosslinked network. Mechanical testing showed that Young’s modulus decreased with increasing PEG content, consistent with the expected plasticizing effect of PEG and the associated increase in chain mobility within the polymer network. By contrast, tensile strength did not vary monotonically with composition, which suggests that fabrication-related factors, such as specimen thickness variation or process-induced defects, also influenced the measured strength. The accelerated acidic degradation data further showed that higher PEG content increased susceptibility to acidic medium penetration, leading to faster mass loss and greater loss of mechanical integrity. The larger reductions observed at early time points may reflect initial removal of more accessible or weakly bound fractions, followed by slower degradation of the remaining crosslinked network. These findings identify PEG content as an effective lever for tuning degradation rate, although higher PEG loading comes at the cost of reduced retention of stiffness and strength. AESO-based materials have previously been studied under alkaline conditions [25]. We show that the same material platform can also degrade in an acidic environment relevant to the stomach while maintaining SLA printability, thermal stability, and initial mechanical integrity. This proof-of-concept study supports continued development of acidic-environment degradable resins for temporary drainage devices, but several translational questions remain open. Future formulation work will need to more precisely tune degradation kinetics, mechanical retention, and long-term functional lifetime, while parallel studies should evaluate biocompatibility, degradation-product safety, and in vivo degradation behavior.

Degradable stent materials may also help address clogging and biofilm-related failure modes. Recent device literature has identified bacterial adhesion, encrustation, and biofilm formation as major complications of indwelling stents [27]. Biodegradable stent concepts have already been paired with anti-biofilm or renewable-surface strategies in the urinary tract [2, 28]. More broadly, our proposed material workflow could be extended to other temporary drainage devices where removal is challenging, such as biliary, esophageal, and ureteral stents, with degradation profiles and mechanics tuned for the target environment.

Additive manufacturing enables fabrication of structurally complex lattice architectures with rapid iteration that is unachievable by conventional manufacturing methods. Current polymeric DPSs are typically manufactured via extrusion from materials such as polyethylene, polyurethane, polytetrafluoroethylene, or proprietary blends of these polymers [29]. These materials and extrusion processes are well-established and economical but geometrically restrictive. In contrast, SLA printing allows access to a unique design space but with limited biocompatible material options. The complex internal structure of TPMS-based stents mandates the use of additive manufacturing. At the same time, 3D printing also enables patient-specific customization with minimal changes to the fabrication workflow. As this framework matures, integrating patient-specific imaging with device-scale simulation and additive manufacturing could enable drainage stents tailored not only in outer dimensions, but also in local lattice density, compliance, and lumen architecture — a translational pathway directly enabled by the design and fabrication approach established here.

TPMS-integrated, additively manufactured stents are a promising design direction for gastric leak drainage. By combining tunable compliance, resistance to kink-induced lumen collapse, preserved or enhanced drainage performance, and a path toward biodegradability, this approach addresses several important limitations of current DPSs and establishes a foundation for future gastric leak-specific drainage devices. The underlying design logic may also be relevant beyond the gastric setting. Biodegradable stents have already been explored clinically and preclinically in the biliary and ureteral settings, where repeat intervention, migration, restenosis, encrustation, or removal remain important challenges [30-33]. These precedents suggest that TPMS-based architected stents could form a broader platform for temporary drainage and luminal-support applications, provided that the geometry and degradation profile are adapted to the anatomical and mechanical requirements of each target location. Taken together, the mechanical, hydraulic, and materials results presented here support the continued development of BRIDGE as a next-generation drainage device.

## Materials and Methods

### TPMS generation in nTop and volume fraction mapping

We generated all TPMS lattice geometries used in this study in nTop engineering design software. The corresponding level-set equations defining each TPMS topology are given below [34]:

Primitive:

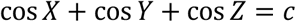

Gyroid:

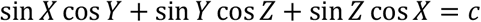

Diamond:

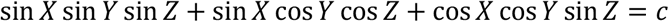

All volume fractions were normalized by the full cylinder volume. Designed volume fraction (VF_D_) denotes the target value prescribed during design and used in the specimen naming convention. Nominal volume fraction (VF_N_) denotes the realized pre-perforation TPMS solid fraction measured in nTop. Effective volume fraction (VF_E_) denotes the realized post-perforation TPMS solid fraction measured in nTop after introducing the central through hole. VF_D_ and VF_N_ differed slightly because nTop requires mid-surface offset (MO), rather than volume fraction, as the design input for TPMS generation. We then estimated the MO corresponding to each target VF_D_ using fitted linear and nonlinear VF—MO relationships (Supplementary Figure 1). Because this translation was approximate, the generated structures showed small deviations between the prescribed VF_D_ and the realized VF_N_. The fitted equations and valid MO ranges for all evaluated topologies and unit cell sizes are summarized in Table 2.

**Table 2.** Equations for VF mapping from mid-surface offsets (MO)

| TPMS topology | UC | Equation ($x = MO$) | $R^2$ | Valid MO range |
| --- | --- | --- | --- | --- |
| Primitive | 0.6 | $y = -26.713x^2 + 6.9792x + 0.5176$ | 1 | -0.05 – 0.10 |
| | 0.7 | $y = -11.872x^2 + 4.9107x + 0.5242$ | 0.9976 | -0.05 – 0.15 |
| | 0.85 | $y = -8.9011x^2 + 4.2026x + 0.5408$ | 0.9979 | -0.05 – 0.15 |
| | 1 | $y = -4.9603x^2 + 3.49x + 0.5212$ | 0.9974 | -0.05 – 0.15 |
| | 1.15 | $y = -3.7762x^2 + 3.0164x + 0.5423$ | 0.9987 | -0.05 – 0.15 |
| | 1.3 | $y = 2.5752x + 0.4482$ | 0.9993 | -0.10 – 0.15 |
| Gyroid | 0.6 | $y = 3.2869x + 0.4982$ | 0.9999 | -0.10 – 0.10 |
| | 0.7 | $y = 2.8537x + 0.4955$ | 0.9997 | -0.15 – 0.15 |
| | 0.85 | $y = 2.3208x + 0.4935$ | 0.9999 | -0.15 – 0.15 |
| | 1 | $y = 1.9601x + 0.5005$ | 1 | -0.15 – 0.15 |
| | 1.15 | $y = 1.687x + 0.5017$ | 1 | -0.15 – 0.15 |
| | 1.3 | $y = 1.4995x + 0.5149$ | 0.9999 | -0.15 – 0.15 |
| Diamond | 0.6 | $y = 3.9144x + 0.5$ | 0.9999 | -0.10 – 0.10 |
| | 0.7 | $y = 3.3549x + 0.5004$ | 1 | -0.10 – 0.10 |
| | 0.85 | $y = 2.7881x + 0.5$ | 1 | -0.10 – 0.15 |
| | 1 | $y = 2.3281x + 0.5$ | 0.9999 | -0.15 – 0.15 |
| | 1.15 | $y = 2.0369x + 0.5017$ | 1 | -0.15 – 0.15 |
| | 1.3 | $y = 1.7675x + 0.5016$ | 1 | -0.15 – 0.15 |

### Finite element analysis for bending load

We performed finite element analysis (FEA) on nTop to evaluate the bending response of tubular triply periodic minimal surface (TPMS) structures. The analyzed TPMS topologies included Gyroid, Primitive, and Diamond, which were generated using the Lattice from Implicit Body domain workflow. All geometries were modeled as skeletal TPMS networks. The non-perforated TPMS cylinder had a diameter of 2.3 mm and a length of 20 mm, whereas the perforated TPMS cylinder had an outer diameter of 2.3 mm, an inner diameter of 1 mm, and a length of 20 mm. Volume fraction (VF) was varied from 0.25 to 1.0, and unit cell size (UC) was varied from 0.6 to 1.3. For all simulations, both ends of the tube were assigned fixed-support boundary conditions, and a downward bending load of 0.01 N was applied at the midpoint of the tube over a 3 mm^2^ area. We solved the models using static analysis solver, which assumes quasi-static loading to equilibrium under a linear load-displacement relationship. The material was modeled as isotropic, linear elastic, with a Young’s modulus of 4.24 MPa and a Poisson’s ratio of 0.3. We determined the Young’s modulus experimentally from tensile testing of dog-bone specimens printed in Flexible 80A resin. The specimens had a width of 5 mm, a thickness of 0.6 mm, and a gauge length of 15 mm. After printing, the samples were post-cured at 60 °C for 20 min and tested in tension at a crosshead rate of 10 mm/min, with three replicates per condition (n = 3). We extracted Young’s modulus from the initial linear region of the stress-strain curve up to 5% strain and used the average value as the input material property for the FEA model. To compare geometric feature size between skeletal TPMS designs, we used nTop’s mesh-thickness workflow to quantify local feature thickness in the final TPMS bodies and summarized the results using average and standard deviation values [35]. This analysis was used to assess the relative fabrication margin between the Gyroid and Diamond topologies.

### Lattice stent design and assembly

We designed BRIDGE devices by combining a central TPMS tube with two pigtail ends, and we used this same overall configuration for all TPMS topologies, unit cell sizes (UC), and volume fractions (VF) evaluated in this study. Each pigtail was modeled in SOLIDWORKS as a curved tube with an inner diameter of 1.68 mm and an outer diameter of 2.3 mm, matching the dimensions of the commercial control stent. The two pigtail ends were positioned with a straight 50 mm gap between them. We then generated a TPMS tubular segment with an outer diameter of 2.3 mm, a length of 50 mm, and a central through hole of 1 mm to form the lattice mid-section, such that the TPMS segment length matched the straight distance between the two pigtail ends. The TPMS segment and the pigtail ends were aligned concentrically and assembled in Materialise Magics. The resulting assembly was verified for overall dimensional consistency with the commercial 7 Fr Advanix™ DPS prior to fabrication.

### TPMS sample fabrication

As a commercial reference, we used a polyethylene Boston Scientific Advanix™ biliary double pigtail stent with a 7 Fr outer diameter and 5 cm length. To generate a digital reference model for comparison with our designs, we manually measured the commercial device using digital calipers and reconstructed its geometry in SOLIDWORKS 2023.

We fabricated printed specimens on a Formlabs Form 3+ stereolithography printer using Flexible 80A resin. All builds used the manufacturer’s Flexible 80A print profile with a layer height of 0.05 mm. The corresponding print settings were 36.0 mJcm^−2^ for perimeter fill exposure, 30.0 mJcm^−2^ for model fill exposure, 144.0 mJcm^−2^ for supports fill exposure, and 72.0 mJcm^−2^ for top surface exposure. Printing was performed in the x-y orientation with the chamber maintained at 35 °C. After printing, all parts underwent solvent cleaning and post-curing to remove uncured resin and complete polymerization, which are critical steps for achieving the final mechanical and biological performance of SLA-printed parts [36]. Printed samples were first washed in isopropanol for 15 min using a Form Wash unit. To further clear residual resin from the internal lumen, we manually irrigated the main channel with 100% isopropanol using a 5 mL syringe fitted with a 24-gauge needle. We then used compressed air to dry the parts and remove any remaining solvent or trapped resin that could obstruct the lumen. The washed parts were subsequently post-cured in a Form Cure unit at 60 °C for 20 min. Finally, we detached the parts from the build platform and manually trimmed the support structures using fine scissors [37]. All fabricated specimens were inspected visually and under stereomicroscopy to confirm lumen patency and absence of gross print defects prior to mechanical or flow testing.

### Tensile test of TPMS geometries

We characterized tensile behavior by uniaxial testing using an Instron 5960 universal testing machine equipped with a 50 N load cell and a tensile gripping fixture. Test specimens consisted of 3D-printed Gyroid tubes fabricated from Flexible 80A resin, with an outer diameter of 2.3 mm, an inner diameter of 1 mm, and a total length of 50 mm. Gyroid designs spanning volume fractions of 0.2–0.5 and unit cell sizes of 0.85–2.0 were tested, with three specimens evaluated per condition (n = 3). Each sample was loaded over a 20 mm gauge length at a crosshead speed of 10 mm/min until failure. Ultimate load was obtained directly from the load-displacement curve, and apparent structural modulus was calculated using the gross cross-sectional area of the tube (1.94 mm^2^).

### Kink curvature test

In the absence of a dedicated standard for double pigtail drainage stents, we adapted flexibility and lumen-patency principles from ISO 25539-2 and FDA guidance no. 1545 to assess kink resistance. These documents use the minimum radius of curvature that a stent can sustain without kinking or exceeding approximately 50% lumen reduction as a benchmark for flexibility evaluation. Based on this framework, we fabricated a 3D-printed conical mandrel with diameters ranging from 30 to 2.5 mm in 2.5 mm increments using Bambu Lab X1 Carbon with PLA basic filament. The height of each step was set to 2.3 mm to match the outer diameter of the commercial and 3D-printed stent samples. Commercial stent and TPMS tubes containing the central through-hole perforation were wrapped around the mandrel beyond 180°, and kink formation was assessed and documented at each diameter. We defined the critical kink condition as the smallest mandrel diameter at which the specimen exhibited no visible kink formation and no major lumen compromise.

### Guidewire-lumen alignment and circumferential material distribution analysis

To examine the effect of guidewire-lumen alignment on perforated TPMS geometry, we generated a Gyroid TPMS block at constant unit-cell size (UC = 1) and designed volume fraction (VF_D_ = 0.3). A tube with an outer diameter of 2.3 mm and an inner diameter of 1 mm was then translated to six different positions within the TPMS block: default (D), center-aligned (C), top-left (TL), top-right (TR), bottom-left (BL), and bottom-right (BR). The intersected geometry was extracted as the final perforated tube. In this way, the relative alignment between the tube lumen and the lattice was varied without altering the underlying TPMS block. For each configuration, the total effective volume fraction (VF_E_) after perforation was measured from the final CAD geometry. To quantify circumferential material distribution, each perforated tube was segmented into 16 equal angular sectors about the lumen center, and the material volume within each sector was calculated. The coefficient of variation of sector-wise volume was then used as a measure of circumferential uniformity, with lower values indicating a more even distribution of material around the lumen. Corresponding TPMS tubes were fabricated in Flexible 80A resin and tested in uniaxial tension (10 mm/min, 50 N load cell, 20 mm gauge length) to determine ultimate load and apparent structural modulus.

### Computational fluid dynamics simulation

To investigate how TPMS architecture affects stent fluid dynamics, we developed a simplified three-dimensional model representing the mid-section of the gastric leak anatomy and the TPMS stent using a representative-volume approach. The model consisted of a TPMS segment measuring 2 mm in length and 2.3 mm in diameter, positioned concentrically at the center of a cylindrical domain with a height of 6 mm and a diameter of 3 mm. We generated the surrounding void space by applying a Boolean subtraction operation in Materialise Magics. Computational fluid dynamics simulations were performed in COMSOL Multiphysics 5.2a. We imposed a pressure inlet of 668 Pa, consistent with typical adult intra-abdominal pressure, and set the outlet pressure to 0 Pa [13]. We modeled the fluid as a Newtonian liquid under no-slip wall conditions and assumed single-phase, incompressible, stationary laminar flow. Because published physical-property data for gastric abscess fluid are lacking, we used a surgeon-informed abscess surrogate and experimentally measured its viscosity (0.137 Pa^·^s) and density (1230 kg/m^3^) for use in benchtop and CFD studies. The model was discretized using a physics-controlled mesh with the “Normal” element size preset. To quantify flow performance, we defined an analysis plane located 0.01 mm downstream of the TPMS face oriented toward the outlet. This plane intersected the full outlet cross section of the geometry, and volumetric flow rate was obtained by surface integration of the velocity field across the plane.

Prior to CFD, the expected flow regime was estimated using channel-scale (Re_ch_) and pore-scale (Re_p_) Reynolds numbers. Following the TPMS hydrodynamic framework previously reported, Re_ch_ and Re_p_ were estimated using the lattice unit outer diameter as a channel length scale and the unit cell size as the pore length scale [16].

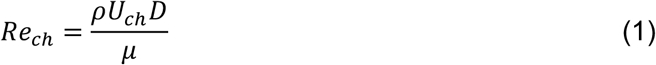

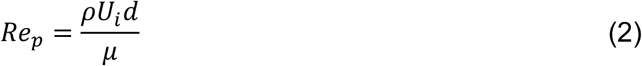

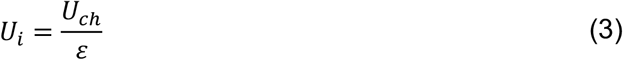

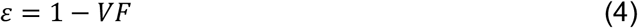

Where *ρ* is the fluid density (1230 kg/m^3^), *U*_*ch*_ is the estimated channel velocity, *U*_*i*_ is an estimated interstitial velocity, *D* is the characteristic channel diameter taken as the TPMS outer diameter (2.3 mm), *d* is the characteristic pore diameter approximated by the unit cell size, *μ* is the fluid dynamic viscosity (0.137 Pa·s), and *ε* is the porosity of the lattice structure. The interstitial velocity was estimated using a standard equation for velocity of fluid in porous media [38]. To estimate the upper bound of *U*_*ch*_, we employed the modified Hagen—Poiseuille’s equation.

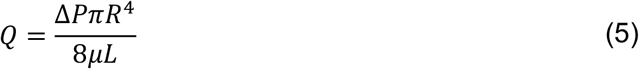

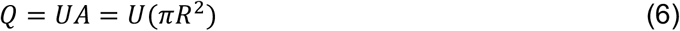

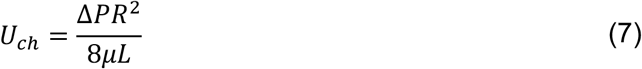

Where Δ*P* is the pressure difference between inlet and outlet, *R* is the radius of the tube, and *L* is the tube length. We assumed an open cylindrical tube of 1.15 mm radius with no obstruction to calculate maximum velocity *U*_*ch,max*_.

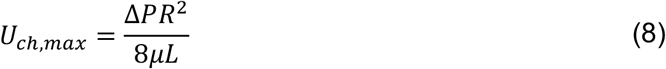

With Δ*P* = 668 *Pa, R* = 1.15 *mm*, and *L* = 2.0 *mm*, we calculated *U*_*ch,max*_ to be 0.4 m/s. Using Eq.8, *Re*_*ch,max*_ was calculated to be 8.32. Additionally, we calculated the *U*_*i*_ for each geometry and the corresponding *Re*_*p,max*_ is provided in Table 3.

**Table 3.** Estimated Reynolds numbers for fluid flowing through TPMS structures.

| Topology | UC (mm) | VF | Porosity $\varepsilon$ | $Re_{p,max}$ |
| --- | --- | --- | --- | --- |
| Primitive | 1.0 | 0.3 | 0.7 | 5.17 |
| Gyroid | 1.0 | 0.3 | 0.7 | 5.17 |
| Diamond | 1.0 | 0.3 | 0.7 | 5.17 |
| Gyroid | 1.0 | 0.2 | 0.8 | 4.53 |
| Gyroid | 1.0 | 0.5 | 0.5 | 7.24 |
| Gyroid | 0.85 | 0.3 | 0.7 | 4.40 |
| Gyroid | 1.5 | 0.3 | 0.7 | 7.76 |

Note that this estimation does not include topology-specific tortuosity effect, hence, Primitive, Gyroid, and Diamond of the same UC and VF have the same estimated *Re*_*p*_. Nevertheless, the range of the Reynolds numbers estimated is well within the laminar flow regime, hence, laminar flow solver was selected.

### Gastric leak benchtop model – fluid flow test

To assess the drainage performance of the stent designs experimentally, we constructed a simplified benchtop platform intended to capture the essential mechanics of endoscopic internal drainage used in the treatment of gastric leaks. The setup was developed with input from bariatric surgeons at Cleveland Clinic Abu Dhabi and was designed to satisfy several functional criteria. We used a water column to impose and sustain a uniform positive pressure around the simulated abscess cavity. Before stent placement, a 3D-printed one-way valve prevented premature outflow of the mock abscess fluid under hydrostatic loading. To improve repeatability across experiments, we designed both the valve and balloon-based abscess module as replaceable components. The simulated abscess cavity consisted of a 12-inch (304.8 mm) transparent latex balloon filled with a viscous test fluid and connected to a soft silicone tube with an outer diameter of 10 mm and an inner diameter of 6 mm, which in turn interfaced with a custom duckbill one-way valve. We designed the valve in SOLIDWORKS (Supplementary Figure 2) and fabricated it using a Stratasys J850 printer, with Agilus30 used for the compliant valve flap and VeroClear used for the rigid housing. To generate the target pressure environment, we submerged the abscess assembly in a plastic container measuring 18 cm in length, 10.9 cm in width, and 10 cm in height and filled it with water to a prescribed height. Based on the hydrostatic pressure relation

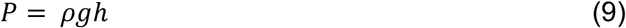

where *P* is pressure, *ρ* is fluid density, *g* is gravitational acceleration, and *h* is fluid height, we set the water level 81.4 mm above the abscess assembly to produce 6 mmHg (800 Pa), consistent with typical adult intra-abdominal pressure [13]. To approximate the viscous contents of an abscess, we consulted three experienced bariatric surgeons, who identified a mixture of 75 wt% concentrated corn syrup in water as a representative fluid. We characterized this surrogate fluid and measured a viscosity of 0.137 Pa^·^s and a density of 1230 kg/m^3^. Each balloon was loaded with 50 mL of the fluid. Assuming complete drainage of the balloon contents, the resulting drop in water level produced a pressure change of less than 0.19 mmHg, indicating that the driving pressure remained effectively constant throughout the test. For each experiment, we placed the abscess assembly in the container and added water to the predetermined height. We then deployed the stent over a guidewire by advancing the guidewire through the silicone tube and duckbill valve until the proximal loop entered the cavity, after which we withdrew the guidewire while keeping the loop fully positioned inside the balloon. During each 20-minute trial, we monitored the mass of drained fluid continuously using a digital balance and recorded the process with a digital camera. We collected data every 5 s for the first 360 s and then every 1 min for the remainder of the experiment. Each stent configuration was tested with three independent replicates (n = 3), and results are reported as mean ± standard deviation.

### Biodegradable resin formulation and printing parameters

We prepared biodegradable SLA resin formulations using acrylated epoxidized soybean oil (AESO, Sigma-Aldrich), polyethylene glycol diacrylate (PEGDA, *M*_*n*_ = 700, Sigma-Aldrich), and polyethylene glycol (PEG, *M*_*n*_ = 200, Alfa Aesar). Four formulations were evaluated with AESO:PEGDA:PEG weight ratios of I (50:50:0), II (45:45:10), III (42.5:42.5:15), and IV (40:40:20). We prepared 20 mL of each formulation for initial screening, except for type III, which was prepared at 200 mL for subsequent print-parameter optimization in a larger resin tank. The formulation procedure was adapted from previous reports on AESO-based photocurable resin systems [25]. AESO, PEGDA, and PEG were combined in a beaker and mixed on a magnetic stirrer hot plate at 50 °C. In parallel, bis(2,4,6-trimethylbenzoyl)-phenylphosphine oxide (BAPO, ionz) was dissolved in acetone at 0.1 g/mL in a tinted glass tube wrapped in aluminum foil to minimize light exposure. After 10 min of mixing, the BAPO solution was added to the resin mixture at 1 wt% of BAPO relative to the total resin mass. The mixture was then stirred at 50 °C for 24 h in the dark to allow complete solvent evaporation. Containers were covered with perforated aluminum foil to permit evaporation while limiting ambient light exposure. To confirm print readiness, 0.1 mL of each resin was dispensed onto a Petri dish and exposed to 405 nm UV light in a Form Cure unit for 5 min, after which formation of a crosslinked solid was visually confirmed.

Resin screening was performed using a customized Formlabs Form 3+ setup that included a 3D-printed tank divider and a CNC-machined aluminum build plate with a custom fixture for printer attachment. Each divided tank compartment measured 50 mm × 60 mm × 25 mm. Cubic and dog-bone specimens were initially printed using the standard Flexible 80A print settings at a layer thickness of 0.050 mm because these geometries did not require fine feature optimization. All four formulations were printed simultaneously using the divided mini-tank configuration under identical conditions, with the printer chamber maintained at 35 °C. Type III was then selected as a representative biodegradable formulation to be used for subsequent print-parameter optimization. After iterative refinement based on observed print quality, the final exposure settings used for type III were 10.0 mJcm^−2^ for perimeter fill, 5.0 mJcm^−2^ for model fill, 115.0 mJcm^−2^ for supports fill, and 80.0 mJcm^−2^ for top surface exposure.

### Accelerated degradation test

We evaluated material degradation using an accelerated degradation assay in simulated gastric fluid. Accelerated aging conditions were selected based on the ASTM F1980 Q10 approach, using a *Q*_10_ value of 2 to approximate the effect of elevated temperature on degradation rate. A *Q*_10_ value of 2 was used because it is a commonly applied default assumption for accelerated aging when material specific degradation kinetics are not readily available. Relative to the physiological reference temperature of 37 °C, incubation at 55 °C corresponds to an accelerated aging factor of 3.48. Thus, 16 days at 55 °C is approximately equivalent to 8 weeks at 37 °C, assuming the degradation mechanism is not altered by the elevated temperature. The degradation medium consisted of 0.2% (w/v) sodium chloride and 0.7% (v/v) hydrochloric acid in water, without added enzyme, and had a pH of 1.0–1.4. Each specimen was immersed in 10 mL of solution in a 15 mL Falcon tube, which was sufficient to fully submerge the sample, and incubated at 55 °C under orbital shaking at 180 rpm. Samples were collected at five time points: day 0, 4, 8, 12, and 16. Day 0 specimens served as untreated controls and were tested as printed without exposure to the degradation medium. For each material type, four specimens were evaluated per time point (n = 4), resulting in a total of 20 samples per formulation. Each specimen was assigned a unique identifier to ensure traceability throughout the study. The degradation solution was not refreshed during the experiment. At each time point, samples were removed from the solution, gently blotted dry without rinsing, and then dried for 24 h in a vacuum desiccator containing silica gel. After drying, we recorded the mass of each specimen and calculated mass change relative to its initial dry mass, after which tensile testing was performed.

## Acknowledgments

We would like to acknowledge the Core Technology Platform (CTP) at New York University Abu Dhabi, specifically Osama Abdullah for technical guidance. This work was also supported by Center for Translational Medical Devices (CENTMED) and Center for Stability, Instability, and Turbulence (SITE).

## Author Contributions

Conceptualization: PP, KBR, JSB, CAV

Methodology: PP, KBR

Investigation: PP, YK, SS, ZM

Technical guidance: OK, MA, WMA

Clinical guidance: JSB, JPP, AZ, CAV, JR, MK

Writing-original draft: PP, KBR

Writing-review and editing: JSB, JPP, OK, MA, AZ, CAV, JR, MK, NM, PN

Supervision: KBR, JSB, CAV

Funding acquisition: KBR, JSB, CAV

## Data and materials availability

All data needed to evaluate the conclusions in the paper are present in the paper and/or the Supplementary Materials.

## Reference

[1] S. A. Firkins and R. Simons-Linares, “Management of leakage and fistulas after bariatric surgery,” Best Practice & Research Clinical Gastroenterology, vol. 70, p. 101926, 2024/06/01/ 2024, doi: 10.1016/j.bpg.2024.101926.

[2] V. L. de Oliveira, A. M. Bestetti, R. P. Trasolini, E. G. H. de Moura, and D. T. H. de Moura, “Choosing the best endoscopic approach for post-bariatric surgical leaks and fistulas: Basic principles and recommendations,” (in eng), World J Gastroenterol, vol. 29, no. 7, pp. 1173–1193, Feb 21 2023, doi: 10.3748/wjg.v29.i7.1173.

[3] I. Siddique, W. Alazmi, and S. K. Al-Sabah, “Endoscopic internal drainage by double pigtail stents in the management of laparoscopic sleeve gastrectomy leaks,” Surgery for Obesity and Related Diseases, vol. 16, no. 7, pp. 831–838, 2020/07/01/ 2020, doi: 10.1016/j.soard.2020.03.028.

[4] P. Woźniewska, I. Diemieszczyk, and H. R. Hady, “Complications associated with laparoscopic sleeve gastrectomy - a review,” (in eng), Prz Gastroenterol, vol. 16, no. 1, pp. 5–9, 2021, doi: 10.5114/pg.2021.104733.

[5] P. Praveenraj et al., “Management of gastric leaks after laparoscopic sleeve gastrectomy for morbid obesity: A tertiary care experience and design of a management algorithm,” (in eng), J Minim Access Surg, vol. 12, no. 4, pp. 342–9, Oct–Dec 2016, doi: 10.4103/0972-9941.181285.

[6] P. Rogalski et al., “Endoscopic management of leaks and fistulas after bariatric surgery: a systematic review and meta-analysis,” (in eng), Surg Endosc, vol. 35, no. 3, pp. 1067–1087, Mar 2021, doi: 10.1007/s00464-020-07471-1.

[7] M. Minakari, V. Sebghatollahi, and M. Namaki, “Success Rate of Endoscopic Treatment for Post-Sleeve Gastrectomy Leaks: A Systematic Review and Meta-Analysis,” Obesity Surgery, vol. 35, no. 8, pp. 3228–3240, 2025/08/01 2025, doi: 10.1007/s11695-025-08014-0.

[8] M. Guirgis, Q. Cheng, D. L. Chan, T. J. Opperman, O. M. Fisher, and M. L. Talbot, “Stage dependent management of sleeve gastrectomy leaks - a systematic review with proposed classification and management algorithm,” Metabolism and Target Organ Damage, vol. 5, no. 3, p. 48, 2025, doi: 10.20517/mtod.2025.10.

[9] L. Su et al., “Modular magnetic microrobot system for robust endoluminal navigation and high–radial force stent delivery in complex ductal anatomy,” Science Advances, vol. 11, no. 43, p. eady4339, 2025, doi: 10.1126/sciadv.ady4339.

[10] C. Chen, Y. Xiong, Z. Li, and Y. Chen, “Flexibility of Biodegradable Polymer Stents with Different Strut Geometries,” (in eng), Materials (Basel), vol. 13, no. 15, Jul 27 2020, doi: 10.3390/ma13153332.

[11] Biliary Devices, Boston Scientific,2015. [Online]. Available: https://www.bostonscientific.com/content/dam/bostonscientific/endo/catalog/Endo_Catalog_2015_Biliary%20Devices.pdf.

[12] C. Brandt-Wunderlich, C. Schwerdt, P. Behrens, N. Grabow, K.-P. Schmitz, and W. Schmidt, “A method to determine the kink resistance of stents and stent delivery systems according to international standards,” Current Directions in Biomedical Engineering, vol. 2, no. 1, pp. 289–292, 2016, doi: 10.1515/cdbme-2016-0064.

[13] B. L. De Keulenaer, J. J. De Waele, B. Powell, and M. L. N. G. Malbrain, “What is normal intra-abdominal pressure and how is it affected by positioning, body mass and positive end-expiratory pressure?,” Intensive Care Medicine, vol. 35, no. 6, pp. 969–976, 2009/06/01 2009, doi: 10.1007/s00134-009-1445-0.

[14] I. Maskery et al., “Insights into the mechanical properties of several triply periodic minimal surface lattice structures made by polymer additive manufacturing,” Polymer, vol. 152, pp. 62–71, 2018/09/12/ 2018, doi: 10.1016/j.polymer.2017.11.049.

[15] X. Zheng, Z. Fu, K. Du, C. Wang, and Y. Yi, “Minimal surface designs for porous materials: from microstructures to mechanical properties,” Journal of Materials Science, vol. 53, no. 14, pp. 10194–10208, 2018/07/01 2018, doi: 10.1007/s10853-018-2285-5.

[16] S. S. Rathore, B. Mehta, P. Kumar, and M. Asfer, “Flow Characterization in Triply Periodic Minimal Surface (TPMS)-Based Porous Geometries: Part 1—Hydrodynamics,” Transport in Porous Media, vol. 146, no. 3, pp. 669–701, 2023/02/01 2023, doi: 10.1007/s11242-022-01880-7.

[17] A. Robinson et al., “Advanced manufacturing of coil-reinforced multilayer vascular grafts to optimize biomechanical performance,” Acta Biomaterialia, vol. 198, pp. 281–290, 2025/05/15/ 2025, doi: 10.1016/j.actbio.2025.04.020.

[18] H. Inam, M. N. Ali, I. R. Jameel, D. Awaiz, and Z. Qureshi, “Development of Robust PEBAX-Based Angiographic Catheter: Design and In Vitro Study,” Materials, vol. 17, no. 17, p. 4248doi: 10.3390/ma17174248.

[19] T.-M. Jung, T. Nairuz, C.-H. Kim, and J.-H. Lee, “Development of a Brain Catheter for Optical Coherence Tomography in Advanced Cerebrovascular Diagnostics,” Biosensors, vol. 15, no. 3, p. 170 doi: 10.3390/bios15030170.

[20] M. Boeykens et al., “Impact of Ureteral Stent Material on Stent-related Symptoms: A Systematic Review of the Literature,” European Urology Open Science, vol. 45, pp. 108–117, 2022/11/01/ 2022, doi: 10.1016/j.euros.2022.09.005.

[21] B. A. Ramesh, J. T. Evans, M. Marietta, and J. Bk, “Suction Drains,” in StatPearls. Treasure Island (FL): StatPearls Publishing Copyright © 2025, StatPearls Publishing LLC., 2025.

[22] M. Nedelcu et al., “Is the Surgical Drainage Mandatory for Leak after Sleeve Gastrectomy?,” Journal of Clinical Medicine, vol. 12, no. 4, p. 1376doi: 10.3390/jcm12041376.

[23] D. Sibilia et al., “Biodegradation Study of Biomaterials Composed of Acrylated Epoxidized Soybean Oil: An In Vitro Study,” BioMed Research International, vol. 2024, no. 1, p. 7100988, 2024/01/01 2024, doi: 10.1155/bmri/7100988.

[24] D. Mondal, A. Srinivasan, P. Comeau, Y.-C. Toh, and T. L. Willett, “Acrylated epoxidized soybean oil/hydroxyapatite-based nanocomposite scaffolds prepared by additive manufacturing for bone tissue engineering,” Materials Science and Engineering: C, vol. 118, p. 111400, 2021/01/01/ 2021, doi: 10.1016/j.msec.2020.111400.

[25] M. Bragaglia et al., “3D printing of biodegradable and self-monitoring SWCNT-loaded biobased resin,” Composites Science and Technology, vol. 243, p. 110253, 2023/10/20/ 2023, doi: 10.1016/j.compscitech.2023.110253.

[26] D. Makharadze, L. J. del Valle, R. Katsarava, and J. Puiggalí, “The Art of PEGylation: From Simple Polymer to Sophisticated Drug Delivery System,” International Journal of Molecular Sciences, vol. 26, no. 7, p. 3102 doi: 10.3390/ijms26073102.

[27] P. Amado et al., “The interplay between bacterial biofilms, encrustation, and wall shear stress in ureteral stents: a review across scales,” Frontiers in Urology, Review vol. Volume 3 - 2023, 2024. [Online]. Available: https://www.frontiersin.org/journals/urology/articles/10.3389/fruro.2023.1335414.

[28] S. Khoddami, B. H. Chew, and D. Lange, “Problems and solutions of stent biofilm and encrustations: A review of literature,” (in eng), Turk J Urol, vol. 46, no. Supp. 1, pp. S11–s18, Nov 2020, doi: 10.5152/tud.2020.20408.

[29] T. H. Baron and J. L. Ponsky, “Chapter 16 - Plastic Pancreatic and Biliary Stents: Concepts and Insertion Techniques,” in ERCP, T. H. Baron, R. Kozarek, and D. L. Carr-Locke Eds. Edinburgh: W.B. Saunders, 2008, pp. 153–163.

[30] G. Song, H. Q. Zhao, Q. Liu, and Z. Fan, “A review on biodegradable biliary stents: materials and future trends,” Bioactive Materials, vol. 17, pp. 488–495, 2022/11/01/ 2022, doi: 10.1016/j.bioactmat.2022.01.017.

[31] Y. Li et al., “Biliary stents for active materials and surface modification: Recent advances and future perspectives,” Bioactive Materials, vol. 42, pp. 587–612, 2024/12/01/ 2024, doi: 10.1016/j.bioactmat.2024.08.031.

[32] Y. Guo, Q. He, M. B. Al-Handawi, T. Chen, P. Naumov, and L. Zhang, “Regulating Supramolecular Assembly and Disassembly of Chitosan toward Efficiently Antibacterial Lubricous and Biodegradable Hydrogel Urinary Catheters,” Advanced Healthcare Materials, vol. 14, no. 6, p. 2404856, 2025/03/01 2025, doi: 10.1002/adhm.202404856.

[33] Y. Wang et al., “A modified biodegradable mesh ureteral stent for treating ureteral stricture disease,” Acta Biomaterialia, vol. 155, pp. 347–358, 2023/01/01/ 2023, doi: 10.1016/j.actbio.2022.11.022.

[34] J. Walles. “What equations are used to create the TPMS types?” nTopSupport. https://support.ntop.com/hc/en-us/articles/360053267814-What-equations-are-used-to-create-the-TPMS-types (accessed.

[35] A. Prasad. “How to measure TPMS wall thickness.” nTopSupport. https://support.ntop.com/hc/en-us/articles/18669009073555-How-to-measure-TPMS-wall-thickness (accessed.

[36] Y. Liu, G. Jin, J.-H. Lim, and J.-E. Kim, “Effects of washing agents on the mechanical and biocompatibility properties of water-washable 3D printing crown and bridge resin,” Scientific Reports, vol. 14, no. 1, p. 9909, 2024/04/30 2024, doi: 10.1038/s41598-024-60450-7.

[37] Formlabs, “Flexible 80A Resin,” 2024. [Online]. Available: https://media.formlabs.com/m/442fcd13220df367/original/-ENUS-Flexible-80A-TDS.pdf

[38] C. W. Fetter, Applied Hydrogeology, 4th ed. Prentice Hall, 2001.

