## Supplementary figures for "Biodegradable Architected Stents for Endoscopic Internal Drainage"

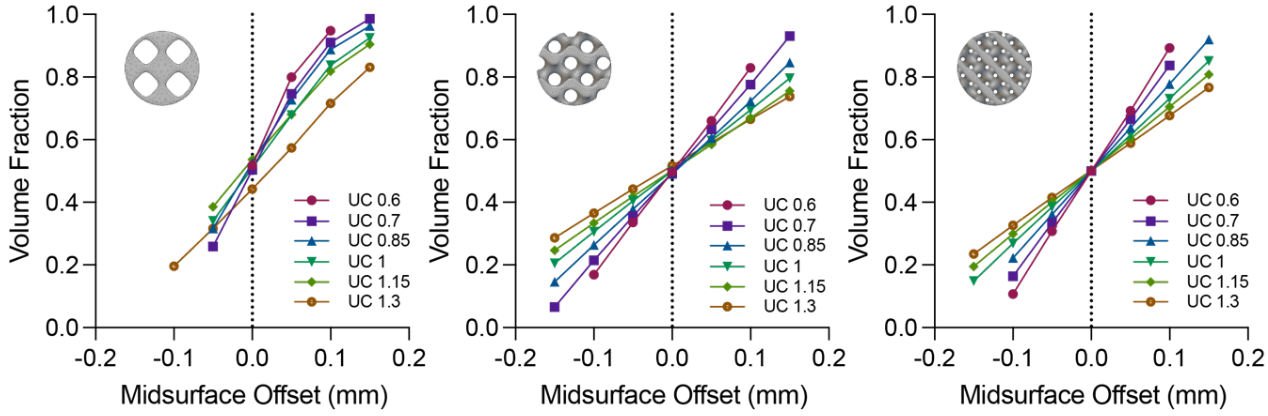


**Supplementary Figure 1.** *Relationship between midsurface offset and volume fraction of TPMS structures generated in nTop.*

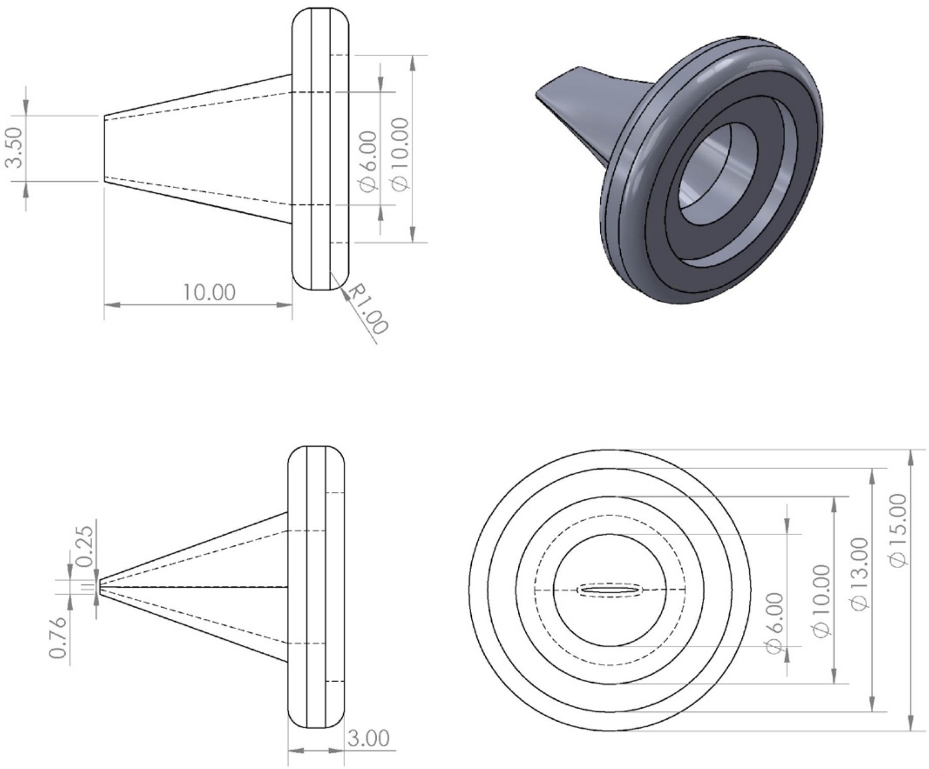


**Supplementary Figure 2.** *Technical drawing with measurement (mm) of the 3D-printed valve on a GL benchtop model.*
